# Serum albumin maintains Wnt water-solubility and activity

**DOI:** 10.1101/2023.01.22.525087

**Authors:** Joo Hye Yeo, Soon Sung Kwon, Jiyeon Chae, Ines Kusen, Jaeeun Han, Ju Yeon Lim, Jae-Ho Cheong, Taeil Kim, Yang Ouk Jung, Chul Hoon Kim, Jinu Lee

**Affiliations:** College of Pharmacy, Yonsei Institute of Pharmaceutical Sciences, Yonsei University, Incheon 21983, Korea; Department of Pharmacology, BK21 PLUS Project for Medical Sciences, Brain Research Institute, Yonsei University College of Medicine, Seoul 03722, Korea; Department of General Surgery, BK21 PLUS Project for Medical Sciences, Yonsei University College of Medicine, Seoul 03722, Korea; Department of Internal Medicine, BK21 PLUS Project for Medical Sciences, Yonsei University College of Medicine, Seoul 03722, Korea; Interpark Bio Convergence Corp., Seoul 05835, Korea; Severance Biomedical Science Institute, Yonsei University College of Medicine, Seoul 03722, Korea

## Abstract

Wnt proteins regulate adult tissue homeostasis and repair by driving stem cell self-renewal and differentiation. High-performance Wnt preparations have enormous therapeutic potential, especially alongside various stem cell technologies. Currently, most of these Wnt preparations contain FBS or the detergent CHAPS to maintain Wnt solubility and activity. Recently, afamin was identified as a serum factor that solubilizes Wnt3a in conditioned media (CM), obviating the requirement for animal sera. Here, we report serum albumin (SA) is required for afamin-mediated solubilization of Wnt3a in CM. Moreover, SA-mediated solubilization of purified Wnt3a in tubes does not require afamin. This means conventional CHAPS-Wnt3a preparations can be modified into SA-purified Wnt3a (SA-pWnt3a) preparations by exchanging CHAPS for SA through dialysis. SA-pWnt3a preparations effectively promote the growth of human stem cell organoids. These data suggest SA as a physiological factor for maintaining Wnt3a activity in therapeutic applications.

## Introduction

Wnt signaling plays important roles in the regulation of a variety of biological processes from development to tissue homeostasis^1^. The activation of Wnt/β-catenin signaling holds enormous therapeutic potential for many disease conditions in which tissue regeneration or the maintenance of tissue homeostasis would be beneficial^2,3^, but there is at least one key hurdle that must be resolved before Wnts can be developed as therapeutic proteins. Wnts are water-insoluble in their active form due to palmitoylation, a post-translational lipid modification^4^. This modification, however, is required for the maintenance of Wnt ligand activity because it facilitates the binding of Wnt to Frizzled (FZD), the receptor for secreted Wnts^5^. Consequently, the formation of an active and soluble Wnt preparation has thus far proven elusive. Additional molecules that help solubilize Wnts in their active acylated forms are necessary. Animal sera (e.g., fetal bovine serum or FBS) can maintain the activity and solubility of Wnt proteins^6^, but the use of animal serum in therapeutic applications can be hazardous for human patients. This is because it can provoke immunological reactions and even lead to pathogen transmission^7-9^. To minimize the contaminants from animal sera that have various biological functions, a method for purifying Wnts in their active form was developed^4^. In this method, Wnt proteins are purified from serum-containing Wnt-conditioned media (CM) using a detergent such as CHAPS during a multi-step column chromatography protocol^4^. The detergent, which remains in the resulting Wnt preparation, can be toxic to cells even at low concentrations^10^. Diluting the CHAPS to a point where it no longer affects cell viability, however, leads to a significant loss of Wnt activity. A Wnt3a-containing CM or a purified Wnt3a, solubilized and stabilized by a clearly defined physiological factor(s), is thus strongly preferred to a detergent-or animal sera-solubilized Wnt3a preparation.

Recently, many groups have become interested in developing a serum-free active Wnt preparation, even for *in vitro* use, due to the increasingly widespread employment of adult stem cell-derived human organoid culture technologies in biomedical research^11^. Since adult stem cell-derived organoid culture techniques were first established with intestinal cells about 10 years ago^12^, Wnt signaling has emerged as the most important *in vivo* niche signaling component for the maintenance of adult stem cells *in vitro*^13,14^. Using this technology, 3D *in vitro* tissue structures that mimic the structural and functional properties of *in vivo* organs can be maintained, passaged, scaled-up, and even stored. Wnt3a was the first member of the Wnts added to culture media to support the long-term *in vitro* proliferation and self-renewal of adult stem cells in organoids^13,15,16^. Typically, this Wnt3a is obtained from the CM produced by a Wnt3a-expressing cell line in the presence of FBS^13^. But unknown components in FBS can have undesirable effects on stem cells, reducing the consistency and performance of this type of Wnt3a formulation^17^. To overcome this limitation, several groups have tried to identify the most critical component(s) in sera that solubilize Wnt and maintain its activity^18,19^. Recently, the Takagi group discovered that afamin (AFM), a member of the albumin superfamily of binding proteins, is a serum component that binds and solubilizes Wnts^19^. By co-expressing Wnt and AFM in the same cell line, soluble but active Wnts can easily be prepared in the absence of animal sera in the form of CM (Wnt3a-AFM CM). Moreover, Wnt3a-AFM CM supports the growth and expansion of human intestinal organoids for more than 18 passages^19^. The structure of AFM has been identified by both glycan and pocket analysis^20^. AFM forms a stable complex with Wnt3a that could improve its solubility by accepting the hydrophobic palmitoleic tail of a Wnt serine residue into a hydrophobic binding pocket on its own surface^20^.

In this study, we found that AFM does not effectively solubilize Wnt3a alone. We have identified serum albumin (SA) as an essential co-factor necessary for Wnt3a solubilization in the production of FBS-free Wnt3a-AFM CM. We also found SA can effectively solubilize and stabilize purified Wnt3a *in vitro* in the absence of AFM. The effective concentration of SA in these two conditions (CM or purified Wnt3a) is far below the normal level present in human plasma or interstitial fluid^21^. In adult stem cell-derived human organoid cultures, both the Wnt3a-AFM CM produced in the presence of SA and the purified active Wnt3a maintained with SA can effectively promote organoid formation. Thus, in this study, we found SA can potently solubilize and stabilize Wnts, making it a simple, cost-effective, and efficient method for producing active Wnt preparations that are more compatible with both *in vitro* and *in vivo* use cases.

## Results

### SA is required for the Wnt ligand activity of CM produced by L-Wnt3a-AFM cells

When the albumin family protein AFM was found to solubilize Wnt3a and maintain its activity in solution^19^, we sought to better uncover the mechanism by which this occurs and determine whether it is influenced by other CM components. To confirm the conclusions reached by Mihara et al.^19^ regarding the role of AFM, we performed a TOPflash assay (Wnt/β-catenin pathway reporter assay) using CM from cells expressing Wnt3a alone (L-Wnt3a cells) or cells co-expressing Wnt3a and AFM (L-Wnt3a-AFM cells) (Figure 1A). Not only did we find significant differences in the amount of Wnt3a in each CM, but the Wnt ligand activities of each CM depended on the presence of AFM (Figure 1B). Thus, we confirmed the reported role of AFM in this Wnt3a CM preparation.

**Figure 1.**
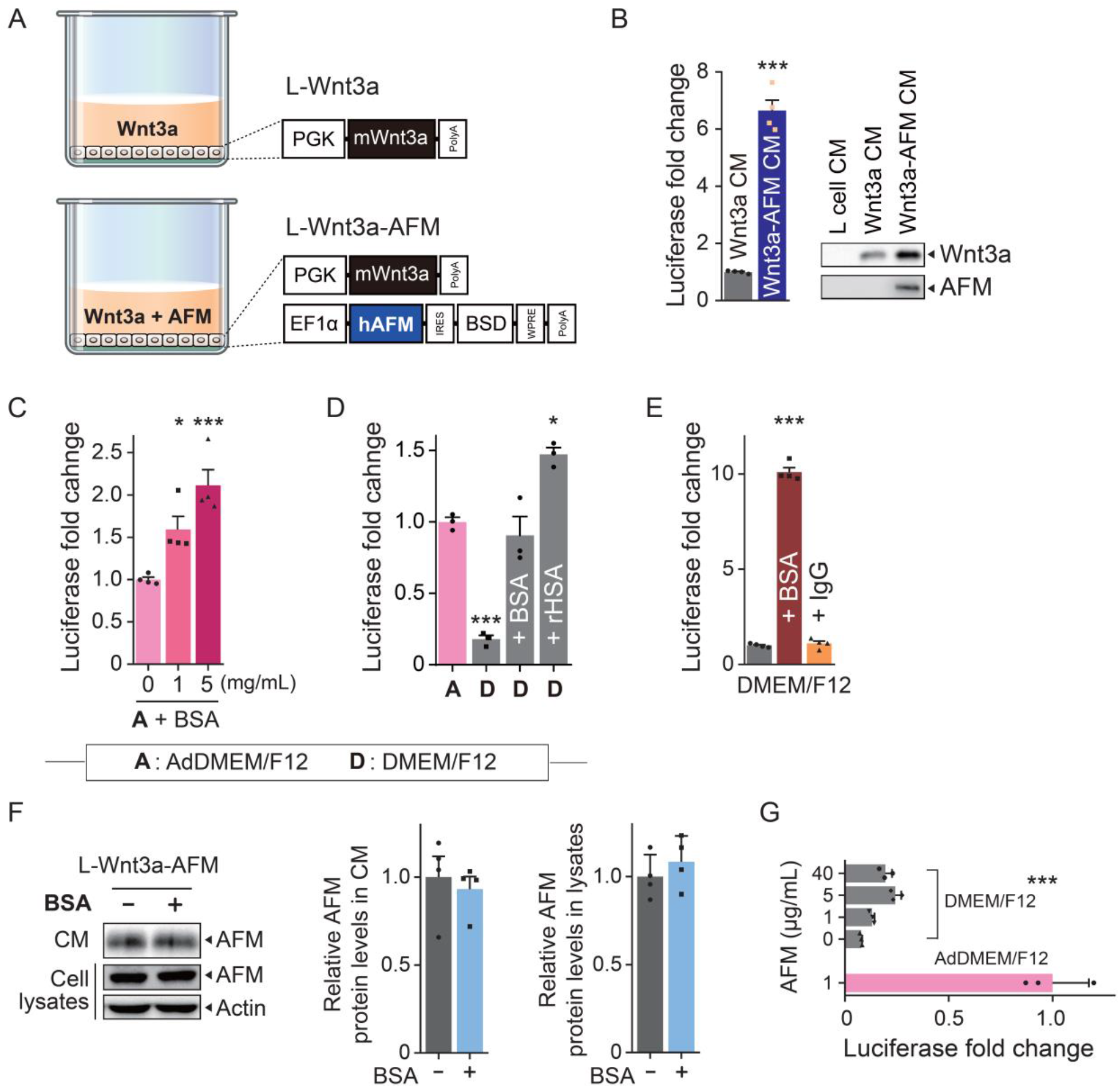
SA is required to maintain the Wnt ligand activity of Wnt3a-AFM CM. (A) L-Wnt3a cells and L-Wnt3a cells transduced with lentivirus harboring pLVX-EF1α-human AFM-IRES-blasticidin-S deaminase (L-Wnt3a-AFM cells) were used to produce Wnt3a CM and Wnt3a-AFM CM, respectively. The media were conditioned by each cell line for 5 days. (B) TOPflash assays (luciferase reporter assays of the Wnt/β-catenin signaling pathway) were performed with both Wnt3a CM and Wnt3a-AFM CM to compare their Wnt3a ligand activities. The luciferase activity with Wnt3a CM stimulation was set to 1 (left). Along with L cell CM, which was used as a negative control, Wnt3a CM and Wnt3a-AFM CM were subjected to anti-Wnt3a and anti-AFM immunoblotting to compare their Wnt3a and AFM levels (right). \*\*\**P* < 0.001, unpaired Student’s *t* test, compared to Wnt3a CM. (C) The effects of increasing amounts of BSA on the Wnt3a ligand activity of Wnt3a-AFM CM were measured via TOPflash. AdDMEM/F12: advanced DMEM/F12. \**P* < 0.05 and \*\*\**P* < 0.001 compared to A + BSA 0 CM; one-way ANOVA followed by post-hoc Bonferroni corrections for multiple comparisons. The luciferase activity when stimulated by Wnt3a CM produced in AdDMEM/F12 was set to 1. (D) TOPflash assays were used to measure the luciferase fold changes of Wnt3a-AFM CM produced in DMEM/F12 compared to Wnt3a-AFM CM produced in AdDMEM/F12. AdDMEM/F12 contains 0.4 mg/ml of BSA, whereas DMEM/F12 lacks BSA. Adding BSA (0.4 mg/mL) or rHSA (0.4 mg/mL) to Wnt3a-AFM CM (in DMEM/F12) restored Wnt3a ligand activity. rHSA: recombinant human serum albumin. The luciferase activity when stimulated by Wnt3a-AFM CM produced in AdDMEM/F12 was set to 1. \**P* < 0.05, \*\*\**P* < 0.001 compared to A (Wnt3a-AFM CM produced in AdDMEM/F12); one-way ANOVA followed by post-hoc Bonferroni corrections for multiple comparisons. (E) Comparison of Wnt3a protein activity resulting from adding either BSA (5 mg/mL) or IgG (5 mg/mL) to Wnt3a-AFM CM (in DMEM/F12). The luciferase activity of Wnt3a-AFM CM (in DMEM/F12) was used as a control and set to 1 on the Y-axis. IgG: Immunoglobulin-G. \*\*\**P* < 0.001 compared to the control; one-way ANOVA followed by post-hoc Bonferroni corrections for multiple comparisons. (F) L-Wnt3a-AFM cells were grown in DMEM/F12 media for 5 days in the presence or absence of 5 mg/mL BSA. AFM protein levels in both CM and cell lysates were measured using immunoblotting (left). Relative protein levels according to their immunoblotting band intensities are indicated in the bar graphs (right). (G) L-Wnt3a cells were grown in DMEM/F12 media for 5 days with varying concentrations of added AFM. TOPflash assays were performed to measure the Wnt ligand activity of Wnt3a (+AFM) CM. The TOPflash activity of Wnt3a CM made in AdDMEM/F12 supplemented with 1 μg/mL AFM was used as a control and set to 1 on the Y-axis. \*\*\**P* < 0.001 compared to the control, one-way ANOVA followed by post-hoc Bonferroni corrections for multiple comparisons. The data (n=3 or 4 biological replicates) are presented as means ± SEM. **Figure supplement 1**. Formulation comparison between DMEM/F12 and AdDMEM/F12 **Figure supplement 2**. MTT assays in L-Wnt3a-AFM cells L-Wnt3a-AFM cells were cultured in the indicated media for 5 days and then MTT assays were performed to measure cell viability or cell proliferation. (Left) BSA (1 or 5 mg/mL) was added to AdDMEM/F12 for 5 days. (Right) L-Wnt3a-AFM cells were cultured in AdDMEM/F12 or DMEM/F12.

Next, we asked whether other components in the CM are required for AFM to produce its effect. We first tested the effect of SA, another member of the albumin superfamily of binding proteins and the most abundant plasma protein, on the Wnt ligand activity of Wnt3a-AFM CM. After adding varying amounts of bovine serum albumin (BSA) to Wnt3a-AFM CM, we measured the resulting TOPflash activity. We found that increases in BSA lead to a proportional rise in Wnt/β-catenin activity, implying that BSA further enhances Wnt3a activity (Figure 1C). We then compared the activity of Wnt3a-AFM CM in two different media: advanced DMEM (AdDMEM)/F12, which contains 0.4 mg/mL BSA and DMEM/F12, which is BSA-deficient (see Figure 1 – Figure supplement 1). As expected, Wnt ligand activity was significantly lower in DMEM/F12 (Figure 1D). Next, we further confirmed the requirement of BSA by showing that the addition of BSA to DMEM/F12-based CM restores its Wnt ligand activity (Figure 1D). This indicates BSA is a potent enhancer of Wnt3a activity in conjunction with AFM. This effect of BSA is unrelated to its potential actions on cell viability or proliferation of L-Wnt3a-AFM cells (see Figure 1- Figure supplement 2). To determine whether this effect of SA is ubiquitous across species, we performed the same test with recombinant human serum albumin (rHSA) derived from rice. We found that, like BSA, rHSA produced an enhancement of Wnt ligand activity (Figure 1D). This result also indicates that the effect of BSA in the TOPflash assay is not due to its contaminating proteins or lipids. We next investigated the specificity of the role of SA in this phenomenon, asking whether other abundant blood-based proteins, such as Immunoglobulin-G (IgG), could give the similar result. After performing a TOPflash assay comparing Wnt3a activity between BSA+ and IgG+ CM, we found that IgG had no effect on Wnt3a activity (Figure 1E). Thus, the phenomenon is specific to BSA.

We next asked whether BSA exerts its effect by increasing the amount of secreted AFM in the CM. We found relatively consistent levels of AFM in the CMs and cell lysates regardless of their BSA status (Figure 1F). This means that BSA has no effect on the amount of AFM synthesized or secreted and that more AFM cannot substitute for the presence of BSA. This was further confirmed when we increased the concentration (up to 40 μg/mL) of exogenous AFM in the absence of BSA and found that it had little effect on Wnt3a activity compared to 1 μg/mL AFM in the presence of AdDMEM/F12 (Figure 1G). Together, these data demonstrate that while AFM in the media is certainly required for Wnt3a activity in CM, so is the presence of BSA.

### SA specifically requires AFM to support the Wnt ligand activity of Wnt3a-AFM CM

After establishing that BSA is necessary for supporting Wnt3a activity in CM, we asked whether higher concentration BSA (5 mg/mL) can maintain Wnt3a activity in Wnt3a CM even in the absence of AFM. To this end, we compared two cell types: L-Wnt3a-AFM cells, which co-secrete Wnt3a and AFM, and L-Wnt3a cells, which secrete Wnt3a alone. When we added exogenous BSA (5 mg/mL) to the DMEM/F12 media in which each cell line was cultured to produce CM, only the resulting Wnt3a-AFM CM showed a dramatic increase in TOPflash activity (Figure 2A). This clear evidence that AFM is essential for BSA to maintain Wnt3a activity in CM led us to further ask whether this capacity is specific to AFM or whether another member of the albumin superfamily of binding proteins, such as vitamin D binding protein (VDBP) or α-fetoprotein (AFP), could take its place. When we compared the TOPflash activities of CMs produced after adding each candidate protein exogenously along with BSA to the cultured L-Wnt3a cells, we found that neither VBDP nor AFP combined with BSA improved Wnt3a activity when compared to BSA alone or BSA combined with exogenous AFM (Figure 2B). These data suggest AFM is the only other member of the albumin superfamily of binding proteins that is effective for supporting Wnt3a activity in combination with BSA in CM.

**Figure 2.**
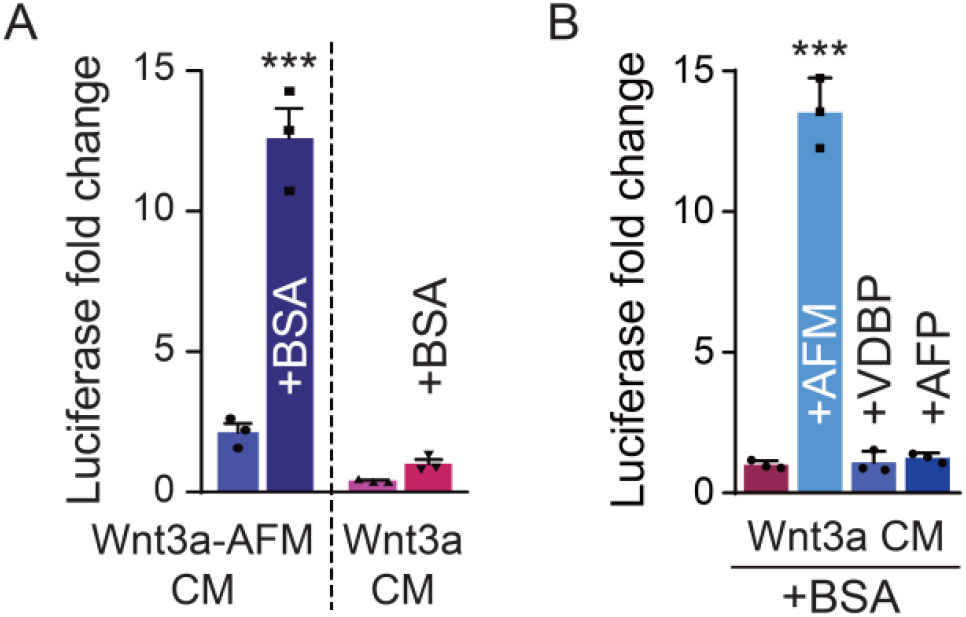
AFM is the only albumin superfamily protein member required for BSA’s maintenance of Wnt3a activity in CM. (A) The effect of exogenous BSA (5 mg/mL) on the Wnt3a ligand activity of Wnt3a-AFM CM (in DMEM/F12) (left) and Wnt3a CM (in DMEM/F12) (right) as measured via TOPflash assays. The luciferase activity when stimulated with Wnt3a (BSA+) CM produced in DMEM/F12 was set to 1. \*\*\**P* < 0.001, unpaired Student’s *t* test, compared to Wnt3a (BSA+) CM. (B) The effect of exogenous co-administration of BSA (5 mg/mL) with AFM (5 μg/mL), VDBP (50 μg/mL), or AFP (10 ng/mL) on the Wnt3a ligand activities of Wnt3a CM (in DMEM/F12). BSA in combination with AFM, VDBP, or AFP was added to the final DMEM/F12 media change for L-Wnt3a cells when producing CM. The resulting CMs were subjected to TOPflash assays to measure their Wnt3a ligand activities. The luciferase activity when stimulated with Wnt3a (BSA+) CM was set to 1 on Y-axis. \*\*\**P* < 0.001 compared to Wnt3a (BSA+) CM; one-way ANOVA followed by post-hoc Bonferroni corrections for multiple comparisons. The data (n=3) are presented as means ± SEM.

### SA prevents aggregation of Wnt3a protein in Wnt3a-AFM CM

Next, we investigated the mechanisms by which SA increases Wnt3a activity in Wnt3a-AFM CM. First, we did not observe any difference in the Wnt3a protein levels of cell lysates from L-Wnt3a-AFM cells cultured in BSA+ or BSA-media (Figure 3A). From these data, we inferred that BSA does not affect the Wnt3a secretory pool. Next, we asked whether BSA affects the secretion of Wnt3a by adding the protein synthesis inhibitor cycloheximide (CHX) to the media of L-Wnt3a-AFM cells and measuring cellular changes in Wnt3a protein levels for up to 24 hrs in the absence or presence of BSA. BSA did not affect the rate of Wnt3a loss in cell lysates (Figure 3B). Rather than changes in Wnt3a production, the reduction in intracellular Wnt3a seems to be due to a change in protein secretion. This is because addition of the Wnt secretion inhibitor IWP-2 produced a near full recovery (Figure 3B). These findings suggest BSA does not affect the typical route by which Wnt3a is secreted. Wnt is normally, at least in part, secreted in exosomes^22^. We found, however, that BSA did not alter the amount of Wnt3a in exosomes derived from Wnt3a-AFM cells (see Figure 3 – Figure supplement 1).

**Figure 3.**
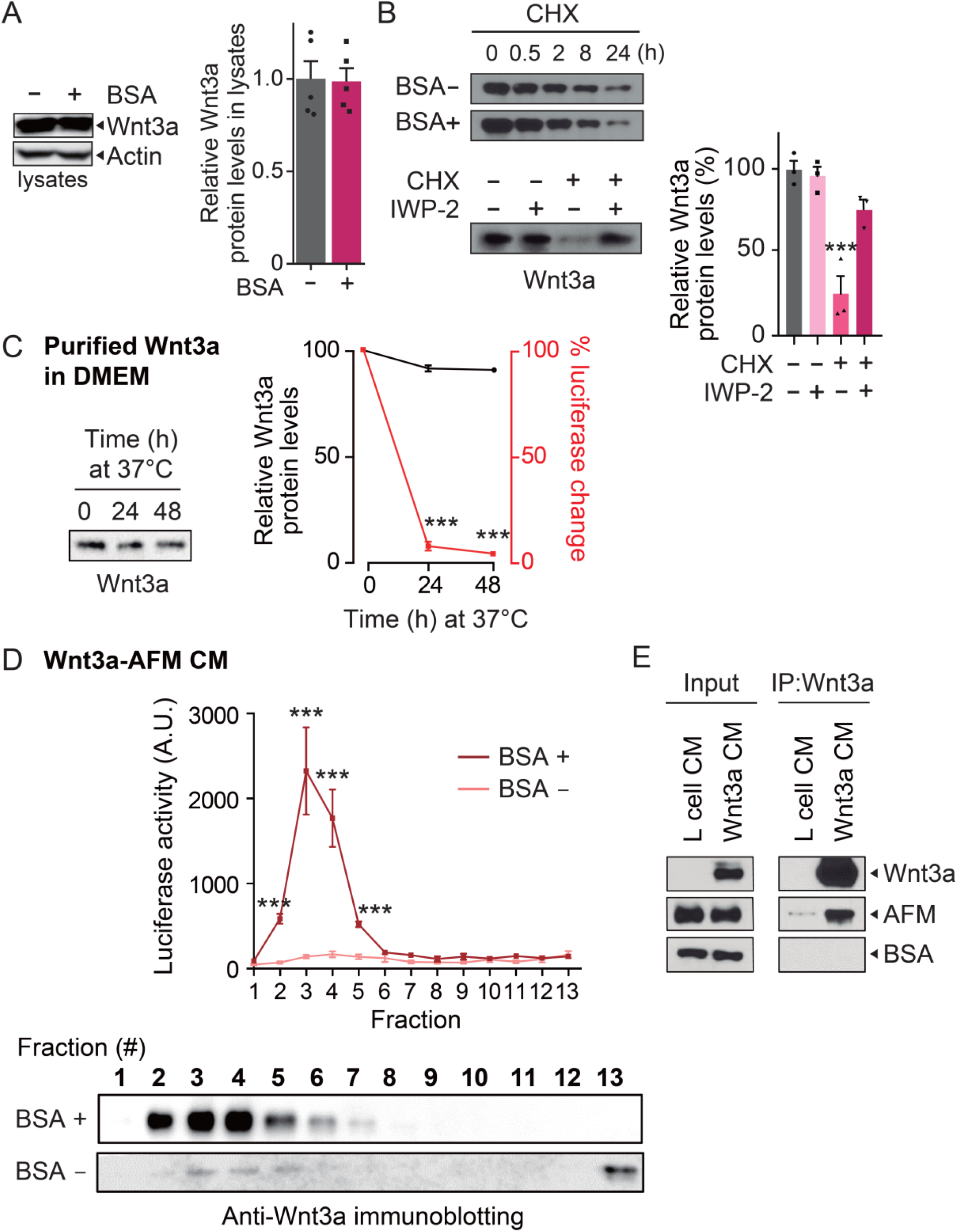
SA solubilizes and stabilizes the Wnt3a-AFM complex. (A) Wnt3a protein levels in the presence or absence of additional BSA (5 mg/mL) were measured via immunoblotting for Wnt3a in cell lysates (left). BSA was added to the media bathing L-Wnt3a-AFM cells for 5 days. The relative Wnt3a protein levels according to immunoblotting band intensities are indicated in bar graphs (right) and compared to the amount of Wnt3a in lysates of L-Wnt3a-AFM cells cultured in DMEM/F12 (BSA-) set to 1. Wnt3a-AFM (BSA-) CM via an unpaired Student’s *t* test. (B) The protein synthesis inhibitor cycloheximide (CHX, 5 μg/mL) was added to L-Wnt3a-AFM cells with or without BSA (5 mg/mL) for the indicated times. Changes in intracellular Wnt3a protein levels over time were monitored using anti-Wnt3a antibodies (upper). In the absence of new protein synthesis, BSA did not affect Wnt3a levels. CHX (5 μg/mL) and IWP-2 (5 μM), which is an inhibitor of Wnt processing and secretion, were added to L-Wnt3a-AFM cells for 15 hrs. The decrease in Wnt3a levels in the presence of CHX can be almost fully attributed due to the secretion of Wnt3a because IWP-2 treatment rescued the loss of Wnt3a protein caused by CHX treatment (lower). \*\*\**P* < 0.001 compared to the CHX-, IWP-2-control, one-way ANOVA followed by post-hoc Bonferroni corrections for multiple comparisons. (C) Purified Wnt3a protein in DMEM/F12 was incubated for the indicated times at 37°C and changes in Wnt3a protein levels were measured via immunoblotting (left). Changes in Wnt3a protein levels and TOPflash activities are depicted in black and red, respectively, in a line graph (right). \*\*\**P* < 0.001 compared to the zero-time control, one-way ANOVA followed by post-hoc Bonferroni corrections for multiple comparisons. (D) Wnt3a-AFM CM produced in the presence and absence of BSA were subjected to sucrose density gradient centrifugation and separated into 13 fractions. The lower numbers indicate the lower-density fractions containing soluble, monomeric or oligomeric Wnt3a; the higher numbers indicate the higher-density fractions containing aggregated, insoluble Wnt3a. Wnt3a ligand activity was measured in each of the 13 fractions using TOPflash assays (upper). The Wnt3a protein level in each fraction was measured via anti-Wnt3a immunoblotting (lower). \*\*\**P* < 0.001 compared to Wnt3a-AFM (BSA-) CM at each fraction, repeated two-way ANOVA followed by post-hoc Bonferroni corrections for multiple comparisons. (E) L cell CM and Wnt3a CM were produced in cultures of L cells and L-Wnt3a cells which were incubated in DMEM/F12 supplemented with AFM 5 μg/mL and BSA 5 mg/mL for immunoprecipitation. L cell CM and Wnt3a CM were subjected to western blotting using anti-Wnt3a, anti-AFM, and anti-albumin antibodies as 2 % input controls. After immunoprecipitating L cell CM and Wnt3a CM with an anti-Wnt3a antibody, the immunoprecipitated proteins were subjected to western blotting using the same sets of antibodies. The data (n = 3–5) are presented as means ± SEM. **Figure Supplement 1**. BSA does not affect Wnt3a secretion on exosomes (A) Exosomes were purified from CM produced in cultures of L-Wnt3a-AFM, L-Wnt3a, or L cells (as a negative control) with or without BSA (5 mg/mL). Differential centrifugation of CM separated it into SN_△_ fractions and P100 exosome fraction. Western blotting for the exosome marker TSG101 confirmed that P100 was enriched with exosomes. Wnt3a was present in P100 pellets of L-Wnt3a or L-Wnt3a-AFM CM (based on DMEM/F12) along with its cargo protein GPR177 (Wntless). BSA did not affect the amount of Wnt3a and GPR177 in exosomes. (B) A quantification of Wnt3a and GPR177 in exosomes is depicted in bar graphs.

We next asked how BSA affects the way Wnt3a’s activity changes after it is released into the CM. Wnt rapidly loses its activity upon its addition to serum- or detergent-free solutions^10,18^. Thus, we looked for any changes in the relative Wnt3a protein level or the TOPflash activity of purified Wnt3a in DMEM/F12 maintained in tubes over 48 hrs at 37°C. Interestingly, although we found TOPflash activity fell rapidly in the absence of BSA, the levels of purified Wnt3a protein did not change over the course of 48 hrs (Figure 3C). Based on this, we hypothesized that BSA supports the long-term maintenance of Wnt3a activity by preventing protein aggregation. To investigate the hypothesis that BSA increases Wnt3a solubility or conformational stability, we performed a protein complex size fractionation, dividing CM with or without BSA into 13 fractions. In the presence of BSA, Wnt3a remained in a monomeric or oligomeric form, with bands present only in the low-density fractions (Figure 3D). In the absence of BSA, however, the Wnt3a bands appeared only in the high-density fractions, indicating the presence of aggregated, inactive Wnt3a. Using TOPflash assays to measure the activity of each fraction, we found the highest Wnt3a activity in the low-density fractions of BSA+ CM containing soluble Wnt3a (Figure 3D). These findings suggest BSA solubilizes and stabilizes active Wnt3a in Wnt3a-AFM CM by preventing its aggregation. AFM is known to form a complex with Wnt3a^19^. To determine whether BSA also binds Wnt3a to prevent its aggregation, we used Wnt3a antibodies to perform an immunoprecipitation of Wnt3a-AFM CM supplemented with BSA (5 mg/mL). We found that while AFM interacts with Wnt3a, BSA does not (Figure 3E). These data suggest a direct interaction between BSA and Wnt3a is not required for BSA to maintain Wnt3a in its active form.

### Wnt3a-AFM CM needs BSA to support the growth and expansion of human adult **stem** cell organoids

We compared the effects of Wnt3a-AFM (+BSA) CM, Wnt3a-AFM (-BSA) CM, and Wnt3a-FBS CM on the growth and passaging of human colon and stomach organoids. For both types of organoids, we found that only Wnt3a-AFM (+BSA) CM, not Wnt3a-AFM (-BSA) CM supported a growth rate comparable to that of Wnt3a-FBS CM (Figure 4A and 4D). Organoids cultured in Wnt3a-AFM (-BSA) CM, which is deficient in BSA, showed a cell growth plateau after the first passage (Figure 4A and 4D). In an immunohistochemistry experiment using the proliferation marker Ki67, we found more proliferating cells in organoids grown in Wnt3a-AFM (+BSA) CM than in those grown in the BSA-deficient CM (Figure 4B and 4E). In a real-time RT-PCR experiment, we also found significantly higher expression of the Wnt target genes *LGR5* and *AXIN2* in both types of organoids in the presence of both BSA and AFM (Figure 4C and 4F). Together, these data indicate BSA is essential for the CM made from L-Wnt3a-AFM cells to support the growth and expansion of adult stem cell organoids. The role of BSA in these preparations seems to maintain Wnt3a in its active form.

**Figure 4.**
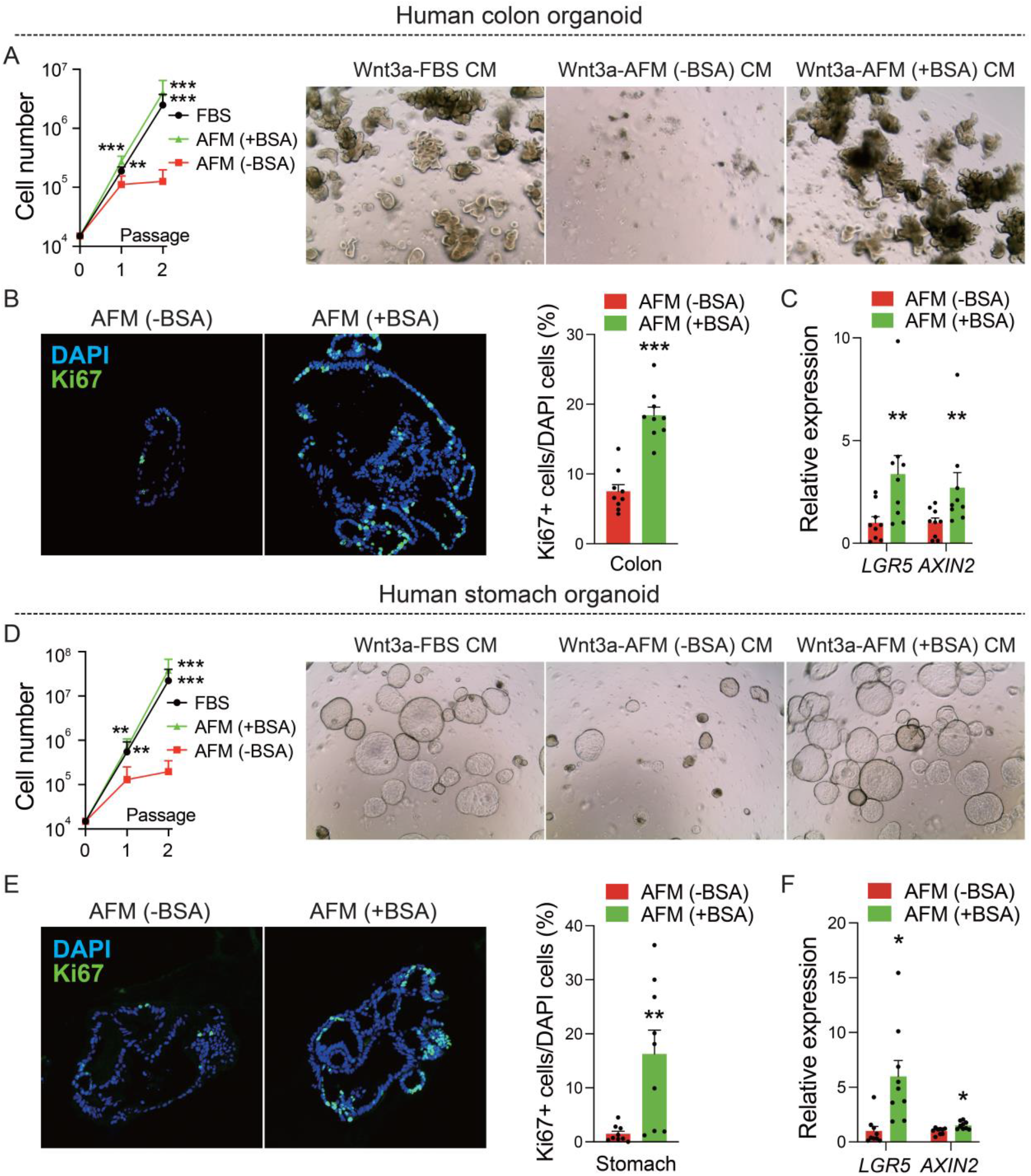
Wnt3a-AFM CM needs BSA to support the growth and expansion of human adult stem cell organoids. (A, D) Growth rate of human colon or stomach organoids in the following conditioned media: Wnt3a-FBS CM, Wnt3a-AFM (-BSA) CM, or Wnt3a-AFM (+BSA) CM. The cells in each well were counted after each passage and the cumulative number of cells at each passage was calculated by the equation in the methods section (left). Image of organoids after 10 days of culture in the indicated conditioned media (right). \*\**P* < 0.01 and \*\*\**P* < 0.001 compared to Wnt3a-AFM (-BSA) CM, repeated measures two-way ANOVA with log-transformed data followed by post-hoc Bonferroni corrections for multiple comparisons. (n=3 biological replicates) (B, E) Human colon or stomach organoids were immuno-stained with the proliferation marker Ki67. Confocal images of fixed organoids processed into paraffin blocks and then immuno-stained with Ki67 (green) and DAPI (blue) (left). The proportion of Ki67+ cells among the DAPI+ cells was calculated and indicated in a graph (right). \*\**P* < 0.01 and \*\*\**P* < 0.001, unpaired Student’s *t* test, compared to Wnt3a-AFM (-BSA) CM. (n=9 biological replicates) (C, F) Quantitative real-time RT-PCR was performed on cDNA synthesized from human colon or stomach organoids cultured in the indicated CM. Expression levels of the Wnt target genes *LGR5* and *AXIN2* were compared. The amount of mRNA (for both *LGR5* and *AXIN2*) when stimulated with Wnt3a-AFM (-BSA) was set to 1 on the Y-axis. \**P* < 0.05 and \*\**P* < 0.01, unpaired Student’s *t* test, compared to Wnt3a-AFM (-BSA) CM. (n=9 biological replicates) The data are presented as means ± SEM.

### SA can solubilize and stabilize purified Wnt3a in the absence of AFM

To directly demonstrate that interactions between Wnt3a, AFM, and BSA are sufficient for the solubilization and stabilization of active Wnt3a, we performed an *in vitro* solubility assay. First, we spun down various combinations of purified Wnt3a with AFM or BSA in solution (incubated for 4 hrs before centrifugation) in a centrifuge to produce a pellet and a supernatant (Figure 5A). We then observed on western blots that Wnt3a was primarily soluble and in the supernatant in the presence of BSA, while it mainly appeared in an insoluble form in the pellet in the absence of BSA (Figure 5B). rHSA also showed a similar effect (see Figure 5 – Figure supplement 1). The addition of AFM did not do much to maintain more of the purified Wnt3a in its soluble form than BSA alone (Figure 5B). Moreover, a fractionation experiment with purified Wnt3a solution demonstrated the occurrence of aggregation even in the presence of AFM (Figure 5C, left panel). It was only in the presence of BSA that Wnt3a was present in its soluble monomeric or oligomeric states (Figure 5C, left panel). We also confirmed the Wnt ligand activity of the soluble Wnt3a via a TOPflash assay of each fraction (Figure 5C, right panel).

**Figure 5.**
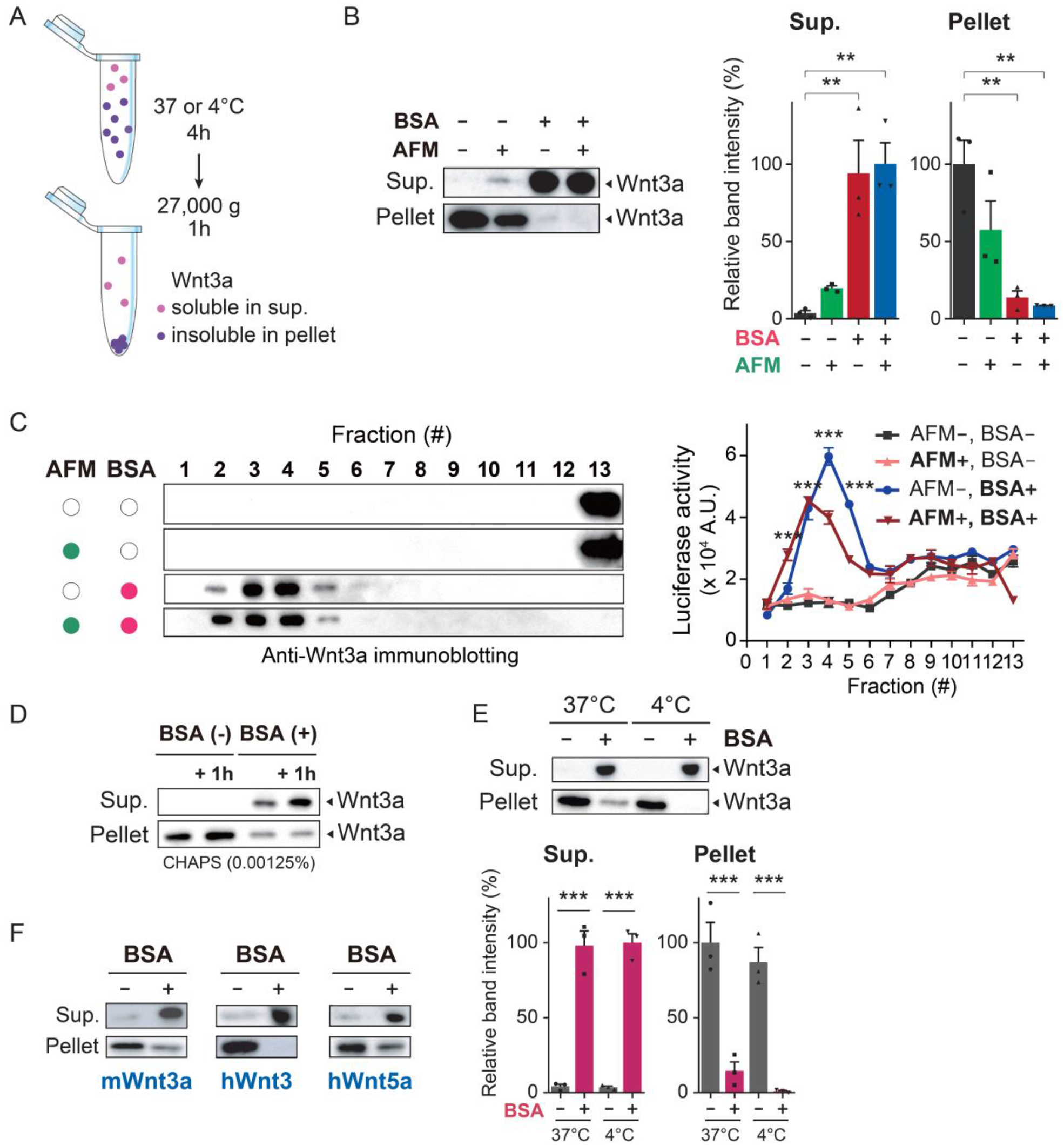
SA solubilizes and stabilizes purified Wnt3a in the absence of AFM. (A) Purified Wnt3a proteins were incubated for 4 hrs in serum-free media (DMEM/F12) containing vehicle, AFM, or BSA at either 4°C or 37°C. Then, the solutions were centrifuged at 27,000 g for 1 hr at 4°C to separate soluble and insoluble Wnt3a. (B) Purified Wnt3a (200 μg/mL with 0.5% CHAPS) was added to a final concentration of 500 ng/mL in DMEM/F12 containing vehicle, BSA (5 mg/mL), or AFM (5 μg/mL). Then, the partitioning of Wnt3a into soluble or insoluble fractions was examined via anti-Wnt3a immunoblotting (left). Relative Wnt3a protein levels in supernatants and pellets are depicted in bar graphs (right). \*\**P* < 0.01 compared to BSA-/AFM-control, one-way ANOVA followed by post-hoc Bonferroni corrections for multiple comparisons. (C) Purified Wnt3a solution was subjected to sucrose density gradient centrifugation as in Figure 3D. The effects of adding BSA and/or AFM were examined. The Wnt3a ligand activity of each fraction was measured via TOPflash assays. \*\*\**P* < 0.001 compared to the BSA-/AFM-control at each fraction, repeated two-way ANOVA followed by post-hoc Bonferroni corrections for multiple comparisons. (D) Purified Wnt3a (200 μg/mL with 0.5% CHAPS) was added to a final concentration of 500 ng/mL in 1 mL of DMEM/F12. BSA was added after aggregating the purified Wnt3a for 4 hrs. The solution was subjected to centrifugation to separate the supernatant and the pellet just after the addition of BSA, or after a 1 hr incubation with BSA. (E) The effect on Wnt3a solubility of adding BSA to purified Wnt3a solution at different temperatures was measured via anti-Wnt3a immunoblotting (upper). Relative Wnt3a protein levels are quantified and depicted in a graph (lower). \*\*\**P* < 0.001 compared to the BSA-control at each temperature, two-way ANOVA followed by post-hoc Bonferroni corrections for multiple comparisons. (F) BSA was added to solutions of multiple members of the Wnt protein family: mouse Wnt3a (mWnt3a), human Wnt3 (hWnt3a), and human Wnt5a (hWnt5a). Protein solubility was examined via a solubility assay as in (A) and the presence of Wnt proteins in the supernatant or in the pellet was determined via anti-Wnt3a immunoblotting. The data (n=3 or 4) are presented as means ± SEM. **Figure supplement 1**. rHSA and BSA show similar effects on Wn3a solubility. Purified Wnt3a (200 μg/mL with 0.5% CHAPS) was added to a final concentration of 500 ng/mL in DMEM/F12 containing vehicle, BSA (5 mg/mL), or rHSA (5 mg/mL). Then, the partitioning of Wnt3a into soluble or insoluble fractions was examined via anti-Wnt3a immunoblotting (left). Relative Wnt3a protein levels in supernatants and pellets are depicted in bar graphs (right). Data (*n* = 3) are presented as means ± SEM. \*\*\**P* < 0.001 compared to BSA-, rHSA-control. **Figure supplement 2**. BSA efficiently solubilizes Wnt3a even in trace amounts of CHAPS Solubility assays were performed with purified Wnt3a co-incubated with different concentrations of CHAPS in the presence or absence of BSA (5 mg/mL). Supernatants (Sup.) and pellets were subjected to western blotting using anti-Wnt3a antibodies.

The purified Wnt3a preparation we used in *in vitro* solubility assays contained trace amounts of CHAPS because we used CHAPS-purified Wnt3a proteins (R&D systems). Wnt3a would become insoluble as the CHAPS was diluted to a concentration at which CHAPS alone could not efficiently prevent Wnt3a aggregation (see Figure 5 – Figure supplement 2). It is notable that, even in trace amounts of CHAPS (0.001 %), BSA was able to maintain Wnt3a in a soluble form without the addition of AFM (see Figure 5 – Figure supplement 2). To further determine whether BSA contributes to the process of solubilizing Wnt3a, we added BSA after Wnt3a aggregation had already occurred. Compared to 1% CHAPS which efficiently solubilized Wnt3a^4^, we found on western blot that a lower concentration of CHAPS (0.001 %) allowed for significant Wnt3a aggregation (Figure 5D). The addition of BSA (5 mg/ml, 4 hrs after Wnt3a aggregation in 0.001 % CHAPS), however, restored Wnt3a solubility even at a lower CHAPS concentration (Figure 5D). This suggests BSA itself can solubilize Wnt3a. We also asked whether BSA affects the thermostability of Wnt3a because a previous study found that purified Wnt3a with CHAPS was unstable at 37°C^23^. We found purified Wnt3a was nearly equally soluble at both 4°C and 37°C in the presence of BSA, indicating that the effect of BSA on Wnt3a activity is unrelated to thermostability (Figure 5E).

Finally, we examined the effects of BSA on other members of the Wnt protein family, seeking to determine the prevalence of its role in protein solubilization. On western blot, we observed a clear shift of mouse Wnt3a, human Wnt3, and human Wnt5a into the supernatant in the presence of BSA (Figure 5F). This suggests multiple members of the Wnt protein family may similarly interact with BSA, presenting opportunities for expanding the methods outlined in this study to other Wnt family proteins. Ultimately, our data indicate BSA effectively maintains purified Wnt3a in its soluble and active form without the help of AFM.

### A new method for preparing purified, active Wnt3a in the absence of CHAPS

Based on the results above, we have developed a new method for maintaining purified Wnt3a in its active form after removing CHAPS. This new method can be used in both *in vitro* organoid culture applications and in the development of safe therapeutic agents. Typically, purified Wnt3a protein preparations contain some CHAPS (0.5–1%), which maintains Wnt3a in an active form. When the CHAPS is diluted, however, Wnt3a loses its solubility and activity (Figure 6A). Because we found that BSA can solubilize and stabilize purified Wnt3a, we planned to dialyze CHAPS from the active Wnt3a solution containing both Wnt3a and BSA (Figure 6B). A purified Wnt3a-BSA solution (pWnt3a-BSA) that remains soluble and active even after the removal of CHAPS would be an ideal Wnt3a preparation. To this end, we placed purified Wnt3a proteins containing CHAPS in a 10 kDa dialysis cassette along with BSA (5 mg/mL) and performed dialysis against standard DMEM/F12 media. Since both Wnt3a and BSA have molecular weights larger than 10 kDa (37.5 kDa and 66.5 kDa, respectively), they remain within the dialysis cup while the 0.6 kDa CHAPS diffuses out into the surrounding media. In contrast to the surface of the low protein binding tubes (LoBind) we used in other experiments of this study, the dialysis membranes adsorbed much of the Wnt3a in the absence of BSA (Figure 6C). Fortunately, the addition of BSA recovered much of this lost Wnt3a. BSA is known to prevent nonspecific protein adsorption to polymer surfaces. Moreover, when we performed a western blot following dialysis, we saw a clear distinction in the partitioning of Wnt3a into the supernatant only in the presence of BSA (Figure 6D). These findings indicate that BSA both maintains protein solubility and prevents the adsorption of Wnt3a to the membrane. These effects seem to be specific to BSA because the same concentration of IgG (5 mg/mL) failed to prevent the adsorption and aggregation of Wnt3a (Figure 6C, D). We also observed that BSA can maintain Wnt3a in its soluble form over time with little change in the amount present in the soluble fraction (Figure 6E, F). To confirm the long-term durability of Wnt3a activity in this preparation, we measured the TOPflash activities of Wnt3a in BSA- and BSA+ solutions stored at 4°C for the indicated number of weeks. We found that purified Wnt3a solubilized and stabilized by BSA remained active for at least 2 wks when stored at 4°C (Figure 6G). We also found that pWnt3a-BSA solution maintained its activity after a freeze-thaw cycle, suggesting mid-to large-scale production of BSA-stabilized Wnt3a and its storage in freezers is feasible (Figure 6H). We also tested the functional validity of the pWnt3a-BSA solution by examining the cooperative interaction between purified Wnt3a and R-spondin. Indeed, we found co-administration of R-spondin and pWnt3a-BSA markedly potentiated canonical Wnt/β-catenin signaling (Figure 6I). This suggests pWnt3a-BSA is compatible with R-spondin, which is critical for the amplification of Wnt signaling^24^. Last, we found pWnt3a-BSA is at least equal to or more efficient than Wnt3a-FBS CM in supporting the growth, expansion, and development of human colon organoids (Figure 6J and 6K). We also confirmed that rHSA similarly supports the growth and expansion of these organoids (see Figure 6 – Figure supplement 1). Together, we present for the first time, a new method for producing purified Wnt3a that remains active in solution, using BSA or even HSA instead of CHAPS as a solubilizing agent.

**Figure 6.**
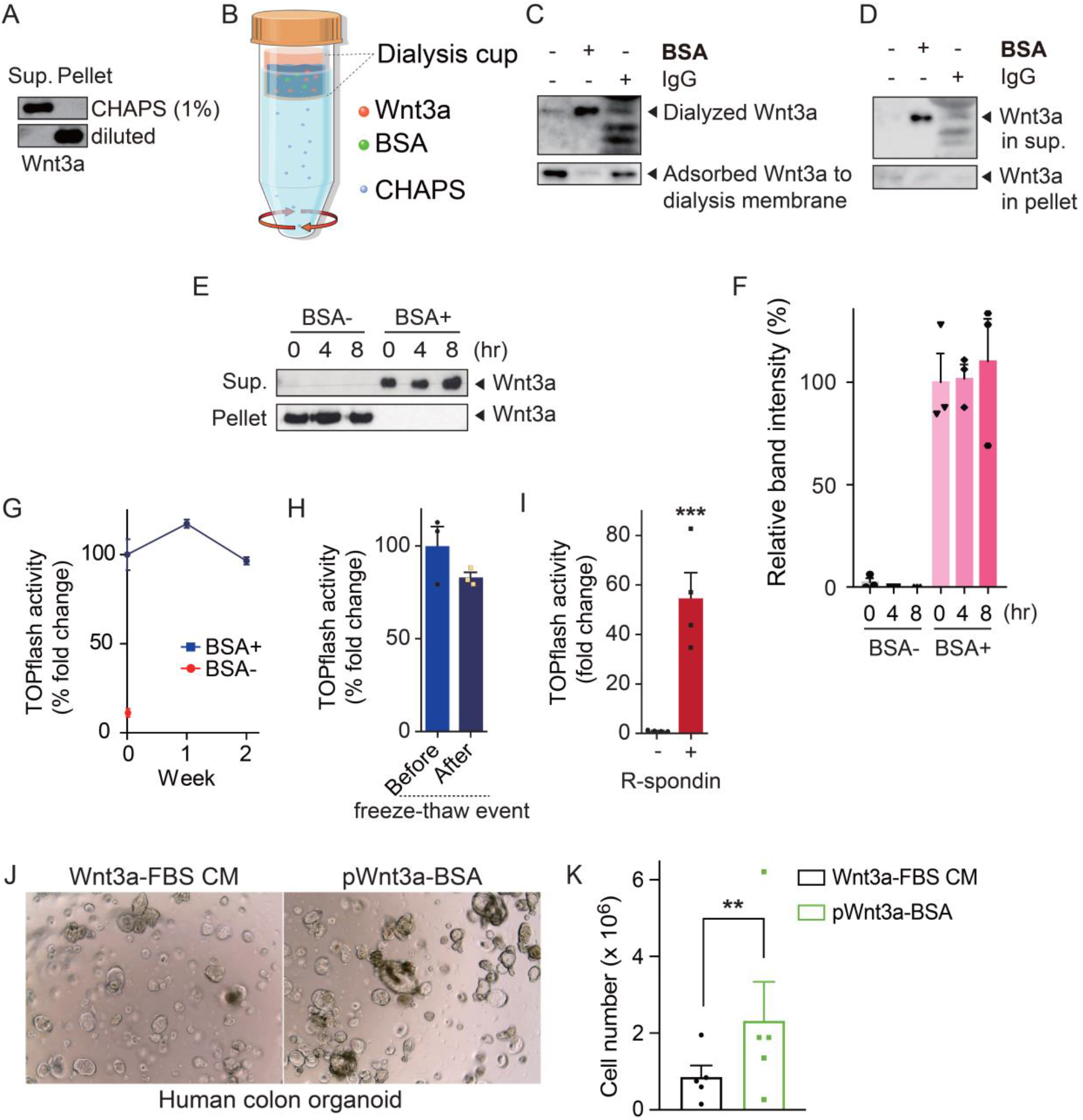
A new method for solubilizing and maintaining purified Wnt3a in its active form in the absence of CHAPS. (A) Adding 1% CHAPS to a purified Wnt3a protein solution maintains Wnt3a solubility. A solubility assay was performed on CHAPS-supplemented (1 %) and CHAPS-diluted (0.001 %) purified Wnt3a solutions as seen in Figure 5A. The protein that remained in the resulting supernatants and pellets was evaluated via anti-Wnt3a immunoblotting. (B) Purified Wnt3a protein (200 μg/mL) containing 0.5% CHAPS was diluted in 2 mL of DMEM/F12 media supplemented with 5 mg/mL BSA. Then, 2 mL of the diluted purified Wnt3a was placed in a 10 kDa cut-off dialysis device. Dialysis was performed against DMEM/F12 media at 4°C for 48 hrs. (C) The effect of adding BSA (5 mg/mL) or IgG (5 mg/mL) to Wnt3a solutions before dialysis on the adsorption of Wnt3a to a dialysis membrane. Wnt3a proteins adsorbed to the dialysis membrane were recovered using 1X Laemmli sample buffer. (D) The effect of adding BSA or IgG to Wnt3a sample solutions before dialysis on their partitioning into supernatants or pellets in a solubility assay of the dialyzed samples. Anti-Wnt3a immunoblotting was performed with supernatants and pellets. (E, F) Purified Wnt3a (200 μg/mL with 0.5% CHAPS) was added to a final concentration of 500 ng/mL to DMEM/F12 containing vehicle or BSA (5 mg/mL) and incubated for the indicated times at 37°C. Then, the partitioning of Wnt3a into soluble or insoluble fractions was examined via anti-Wnt3a immunoblotting. Relative Wnt3a protein levels in supernatants are depicted in bar graphs. The band intensity of BSA+ 0 hr was set to 100. \*\*\**P* < 0.001, one-way ANOVA followed by post-hoc Bonferroni corrections for multiple comparisons, compared to corresponding BSA-time control. (G) Changes in TOPflash activities of purified Wnt3a dialyzed in the presence or absence of BSA (5 mg/mL), after being stored at 4°C for indicated wks, were monitored for 2 wks. (H) Activities of dialyzed purified Wnt3a-BSA solutions (pWnt3a-BSA, 1 μg/mL) were tested via TOPflash assays after a freeze-thaw cycle. (I) Wnt ligand activities of pWnt3a-BSA solutions (1 μg/mL) measured via TOPflash assays after adding R-spondin1. The luciferase activity when stimulated with pWnt3a-BSA solution was set to 1. \*\*\**P* < 0.001, unpaired Student’s *t* test, compared to solutions without R-spondin1. (J) Images of human colon organoids after 10 days of culture in indicated media. (K) Growth rate of human colon organoids in indicated media. Cell numbers of each well were counted and the cumulative number of cells after passage 2 was calculated by the equation mentioned in the method. \*\**P* < 0.01, ratio paired *t* test compared to Wnt3a-FBS CM. Data (n=3-5 biological replicates) are presented as means ± SEM. **Figure supplement 1**. rHSA and BSA show similar effects on growth and expansion of human colon organoids Growth rate of human colon organoids in indicated media (Wnt3a-FBS CM or pWnt3a-rHSA). Cell numbers of each well were counted and the cumulative number of cells after passage 2 was calculated by the equation mentioned in the Method. \*\**P* < 0.01, ratio paired *t* test compared to Wnt3a-FBS CM. Data (*n* = 5 biological replicates) are presented as means ± SEM.

## Discussion

Mihara et al. (2016) found AFM binds Wnt3a in a 1:1 ratio in the CM from Expi293F cells co-expressing AFM and Wnt3a, maintaining Wnt3a in a soluble state in CM. In this study, our recognition of a difference between the ability of basal media such as DMEM/F12 and AdDMEM/F12 to solubilize the Wnt3a-AFM complex led us to identify SA as an additional important factor in serum that can improve the yield and performance of active Wnt3a CM preparations. This does not mean AFM is not essential in making serum-free Wnt3a CM. Rather, we found that although SA alone has minimal effects on solubilization or stabilization of Wnt3a in Wnt3a CM production, we saw a large increase in Wnt3a activity with the addition of both BSA and AFM. From their data, we inferred that Mihara et al.^19^ likely used AdDMEM/F12 factory-formulated to contain 0.4 mg/mL of BSA. They showed on a protein gel stained with Oriole fluorescent stain that purified Wnt3a-AFM from the eluted Wnt3a-AFM CM fractions was contaminated by BSA. We think this covertly contributed to their identification of AFM as the serum factor that solubilizes and protects Wnt3a. After comparing the activity of all the members of the albumin superfamily proteins, including SA, AFM, α-fetoprotein, and vitamin D binding protein, Mihara et al.^19^ concluded that SA is not involved in maintaining active Wnt3a. This finding is consistent with our data showing that BSA alone cannot efficiently solubilize Wnt3a in the process of CM production. Because SA indeed requires AFM to maintain active Wnt3a secreted from cells, it is highly possible AFM has a distinct role from BSA. Naschberger et al.^20^ revealed how afamin forms a stable complex with Wnt proteins and Mihara et al.^19^ showed that Wnt3a is nearly absent in CM when AFM is removed. However, AFM does not effectively stabilize purified Wnt3a in tubes^10^. Considering these results, the role of AFM may require complex formation between AFM and Wnt3a during the process of its synthesis, transport, or secretion. In contrast, according to our results and consistent with a previous report^19^, SA does not affect Wnt3a secretion. Further studies on possible mechanisms by which AFM functions in the Wnt secretory pathway may clarify the well-known Porcupine and Wntless-mediated Wnt secretion pathway.

A previous study using Wnt-responsive luciferase reporter cells found that heparan sulfate proteoglycan (HSPG) rather than BSA stabilized purified Wnt3a protein activity^18^. This result is inconsistent with our current data showing that BSA can solubilize and stabilize purified Wnt3a. We found that pWnt3a-BSA efficiently increases Wnt/β-catenin pathway reporter expression and promotes the growth of adult stem cell organoids. The discrepancy appears to come from the concentrations of BSA tested in each study. We used 0.4 mg/mL or 5 mg/mL BSA, whereas the previous study only tested concentrations up to 30 μg/mL. Considering the typical concentrations of BSA in the blood (30–50 mg/mL) and interstitial fluids (around 30% of that in the plasma), the concentrations of BSA we used are more physiologically relevant and are unlikely to cause any cytotoxicity.

The structural modeling of the albumin-Wnt3a complex has not yet been successful. This is because the albumin binding pocket, although deep and branched, is narrower and its entrance region presents charged and polar residues^20^. This discourages the entry of hydrophobic palmitoleic O-acylation moieties, resulting in a much-reduced probability of direct interaction between SA and Wnt3a. Our immunoprecipitation results also indicates that SA does not participate in any protein-protein interactions with Wnt3a the way AFM dose. Instead, SA may work indirectly by protecting afamin from taking forms with lower Wnt3a binding affinities. This is unlikely, however, because purified Wnt3a can be solubilized and stabilized by SA alone, even in the absence of AFM. It is also unlikely that BSA binds AFM directly because there is a clear separation of SA from the Wnt3a-AFM complex when they are run through a size exclusion chromatography column^19^. On the molecular level, albumin does not have a strong chaperone-like activity^25^. It is possible that Wnt3a stabilization occurs via weak nonspecific interactions with albumin that become physiologically relevant at higher albumin concentrations. Further studies will be necessary to clarify the exact mechanism by which SA solubilizes Wnt3a and whether its function applies only to the Wnts.

Wnt preparations hold promise as additives to the media used for stem cell cultures, especially adult stem cell-derived human organoid cultures^15^. Such organoids have enormous diagnostic and therapeutic potential in precision medicine^11^. Our identification of the most abundant serum protein (35–50 mg/mL)^21^ as a critical factor that supports the prolonged maintenance of active Wnts is encouraging because it provides a solubilization and stabilization strategy that is both physiological and robust. SA has a long half-life of 21 days^26^, suggesting that it will prove to be a durable Wnt stabilizer. Recombinant human SA produced in rice gives an equivalent, or even superior, performance in stabilizing the Wnt3a-AFM complex. This will make it even easier to produce safe Wnt3a at the high yields. We expect the use of purified Wnt3a preparations to be far preferable to CM in organoid cultures. pWnt3a-BSA has less animal cell-derived contamination, which is beneficial when culturing cells as sensitive as stem cells. It is also now possible to add measurable amounts of Wnt3a into cultures with minimal variability, permitting more precise comparisons of stem cell and organoid culture results from different laboratories. We have not yet tested the long-term effects of pWnt3a-BSA on organoid cultures, but it seems likely that even more advantages can be found, such as the maintenance of stemness and improvements to differentiation or maturation potential.

SA is cheaper and seems to act via a more “physiological” mechanism than other currently known Wnt stabilizers, such as heparan sulfate proteoglycan (HSPG), secreted Frizzled-related protein (sFRP), and liposomes^10,18,27^. Although HSPG is also a physiological component of FBS, it is too expensive for producing HSPG-stabilized Wnt3a preparations of practical use. sFRP both promotes and suppresses Wnt/β-catenin signaling, depending on the context^28^. Liposomes mixed with purified Wnt3a are an effective stabilizer of Wnt3a^10^, but they are costly to produce and it is technically demanding to prepare liposomal Wnt in standard biology labs. Moreover, liposomes sometimes undergo leakage of their encapsulated molecules and suffer from phospholipid oxidation, limiting their effectiveness. Wnt3a preparations stabilized with SA overcome all the drawbacks of these other Wnt stabilizers. And perhaps most important, the SA-based preparation of Wnt3a we propose here is simpler to make and more physiological, seemingly mimicking how human organs stabilize their extracellular Wnts.

In this study, we identified SA as an essential and physiological factor that maintain Wnt3a in its soluble and active form. The use of SA in preparing Wnt3a-containing CM or purified Wnt3a will greatly contribute to higher-yield production and purification of active Wnt3a and other Wnts. This will pave the way for the optimization of Wnt preparations for various therapeutic uses and as a culture additive in producing high-quality adult stem cell-derived human organoids.

### Experimental procedures

#### Reagents and consumables

Bovine serum albumin (BSA), recombinant human albumin (rhALB), and immunoglobulin G (IgG) were purchased from Sigma-Aldrich (St. Louis, MO, USA). Protein LoBind tubes were purchased from Eppendorf (Hamburg, Germany). Human Wnt3a, Mouse Wnt3a and human Wnt5a proteins were purchased from R&D systems (Minneapolis, MN, USA). Human Wnt3 and afamin protein were purchased from Origene (Rockville, MD, USA) and R&D systems, respectively.

### Cell culture

HEK293 STF (ATCC, CRL-3249), Wnt3a-producing L cells (L-Wnt3a cells), and L-Wnt3a cells transduced with pLVX-EF1α-human afamin-IRES-blasticidin-S deaminase (BSD) (L-Wnt3a-AFM cells) were grown in Dulbecco’s Modified Eagle’s medium (DMEM, Hyclone) supplemented with 100 IU/mL penicillin, 100 μg/mL streptomycin, and 10% fetal bovine serum (FBS, Hyclone). Cells were maintained in a 5% CO_2_/95% air incubator with a humidified environment at 37°C. The L-Wnt3a cells were a kind gift from Hans Clevers.

### Plasmid construction

To generate the lentiviral human AFM expression plasmid, pLVX-EF1α-IRES-BSD was first generated by modifying pLVX-EF1α-IRES-Puro (Clontech). Then, the human AFM CDS was amplified by PCR from pCR-BluntII-TOPO-AFM (MHS6278-211689548; Dharmacon, Lafayette, CO, USA) with the primers listed in Table 1. The product was digested with XbaI and inserted into the XbaI site of pLVX-EF1α-IRES-BSD. Primer used for constructing the AFM expression plasmid is as described below.

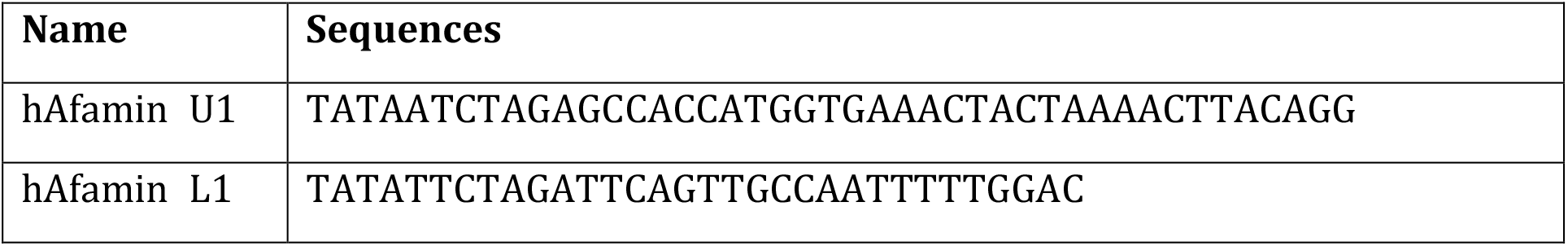

### Generation of L-Wnt3a-AFM cells by lentiviral transduction

HEK293T cells (human embryonic kidney cells) were seeded on 6-well plates pre-coated with poly-L-lysine (PLL) at a density of 600,000 cells/well and incubated for 24 hrs. The cells were transfected with 3 μg of a 4:3:1 mixture of lentiviral human AFM expression plasmid, packaging plasmid (psPAX2, Addgene 12260), and envelope expression plasmid (pMD2.G, Addgene 12259) in the presence of polyethyleneimine. The cells were refreshed with 3 mL of growth medium 16 hrs after transfection and further incubated for 36 hrs. Then, the media containing the lentiviruses were harvested and centrifuged at 3,000 rpm for 3 min to eliminate cell debris. The supernatants were collected and stored at −80 °C until use.

One day before transduction, L-Wnt3a cells were plated on 24-well plates at a density of 50,000 cells/well and grown for 24 hrs. The culture medium was replaced with 400 μL of a 1:1 mixture of the lentivirus-containing medium and fresh culture medium supplemented with polybrene at a final concentration of 4 μg/mL. After 15 hrs, the cells were refreshed with 500 μL of culture medium and incubated for an additional 72 hrs. L-Wnt3a-AFM cells were generated by selection with 2 μg/mL puromycin until a parallel culture of L-Wnt3a cells died.

### Production of Wnt CM

L-Wnt3a-AFM or L-Wnt3a cells were plated at a density of 150,000 cells per 24-well plate or 4,500,000 cells per 100-mm plate and incubated for 48 hrs. Then, the cells were washed twice with phosphate-buffered saline (PBS), refreshed with the indicated medium (500 μL for the 24-well plates; 12 mL for the 100-mm plates), and incubated for 5 days. The CM was harvested and centrifuged at 1,000 rpm for 2 min. The supernatants were cleared through a 0.45 μm syringe filter and stored at 4 °C.

### Luciferase reporter assay to measure Wnt activity

HEK293 STF cells were seeded on PLL-coated 96-well opaque plates at a density of 25,000 cells/well and grown for 24 hrs. Then, each well was treated with 100 μL of fresh culture medium combined with 50 μL of CM to assay for 24 hrs. After aspiration, 100 μL of the 1x steady-Glo reagent (Steady-Glo® Luciferase Assay System, E2520; Promega, Madison, WI, USA) was added to each well. The luminescence of each well was measured using an Infinite 200 PRO plate reader (Tecan).

### MTT assay for assessing cell viability

The growth medium was removed from the cells and 1 mg/mL 3-(4,5-Dimethylthiazol-2-yl)-2,5-diphenyltetrazolium bromide (MTT) dissolved in growth medium was added to each well of interest. The cells with the MTT solution were incubated in 5% CO_2_/95% air at 37°C for 2 hrs. Then, after aspirating the MTT solution, solvent solutions (10% sodium dodecyl sulphate and 25% dimethyl formamide, pH 4.7) were added to each well and incubated overnight. The solutions were then transferred to a 96-well plate and the absorbance at 590 nm was measured.

### Immunoblot

CM samples were harvested and centrifuged at 1,000 rpm for 2 min. Then, the supernatants were mixed with 5× Laemmli sample buffer and boiled for 10 min. For cell lysate samples, the cells were placed on ice and washed twice with cold PBS containing 100 mg/L CaCl_2_ and 100 mg/L MgCl_2_. The cells were lysed with cold RIPA lysis buffer containing 10 mM Tris-Cl (pH 8.0), 1% Triton X-100, 140 mM NaCl, 1 mM ethylenediaminetetraacetic acid (EDTA), 0.1% sodium dodecyl sulfate, and 1× cOmplete Protease Inhibitor Cocktail (Roche, Basel, Switzerland). The lysates were clarified by centrifugation at 13,000 rpm at 4 °C for 10 min. The protein concentration was measured using a bicinchoninic acid (BCA) assay. Samples with identical concentrations were added to 5× Laemmli sample buffer and boiled for 10 min. For purified Wnt proteins, samples were mixed with 5x Laemmli sample buffer and boiled for 10 min. The denatured samples in Laemmli buffer were separated by 12% sodium dodecyl sulfate polyacrylamide gel electrophoresis (SDS-PAGE) and transferred onto nitrocellulose membranes (Whatman, Dassel, Germany). The membranes were blocked with 5% skim milk in PBS containing 0.05% Tween 20 (PBST). Primary antibodies (anti-mouse Wnt3a [2391; Cell Signaling Technology, Danvers, MA, USA], anti-human AFM [sc-373849; Santa Cruz Biotechnology, Santa Cruz, CA, USA], anti-human albumin [ab207327; Abcam, Cambridge, UK] and anti-β-actin Ab [sc-1651; Santa Cruz Biotechnology]) were used at a 1:1,000 dilution in blocking solution. Secondary antibody conjugated to horseradish peroxidase was used at a 1:5,000 dilution in skim milk. Immunoblot images generated with enhanced chemiluminescence (ECL) solution (Thermo Fisher Scientific) were captured using a Fusion Solo 4M (Vilber Lourmat, Eberhardzell, Germany) camera. The band intensities were analyzed using ImageJ software.

### Evaluation of Wnt stability

The indicated Wnt3a CM (Wnt3a-AFM CM with or without BSA, Wnt3a-FBS CM) or purified Wnt3a was added to Protein LoBind tubes and further incubated for 6, 12, 24, or 48 hrs in 5% CO_2_/95% air at 37°C. Immunoblotting with anti-Wnt3a antibodies and TOPflash assays were conducted to assess Wnt3a protein levels and Wnt3a ligand activities at the indicated time of incubation.

#### Sucrose density gradient centrifugation

Sucrose density gradient centrifugation was conducted as previously described, with minor modifications. One milliliter each of 20%, 15%, 10%, and 5% sucrose solutions was layered in a 5 mL polyallomer tube (326819, Beckman Coulter) from bottom to top. Then, 1 mL of the indicated Wnt preparation (Wnt3a CM or purified Wnt3a) was loaded on the top of the sucrose gradient. The tubes with their samples were centrifuged at 150,000 x g at 4°C for 5 hrs. The centrifugation was performed with an Optima MAX-XP ultracentrifuge using an MLS 50 rotor (Beckman Coulter). Then, 400-μL fractions were collected from top to bottom and stored at 4°C. Each fraction was analyzed for Wnt activity via TOPflash assays and for Wnt protein amount via immunoblotting using anti-Wnt3a antibodies.

#### Solubility assays on purified Wnt

Solubility assays were conducted as previously described^29^, with minor modifications. Wnt proteins (500 ng/mL) were incubated in serum-free media containing vehicle, AFM, or BSA at 37°C. After incubation for the indicated time, the solutions were centrifuged at 27,000 g for 1 hr at 4°C. The supernatants were collected and mixed with 5× Laemmli sample buffer, and the pellets were dissolved with 1× Laemmli sample buffer. The samples were then subjected to immunoblotting, and the amount of Wnt protein in each fraction was detected using Wnt subtype-specific antibodies.

#### Exosome purification

Exosomes were purified by differential centrifugation as previously described ^22^. Briefly, Wnt3a CMs harvested from L cells or L-Wnt3a-AFM cells with or without BSA were subjected to sequential centrifugation steps of 300 g, 2,000 g, and 10,000 g for 10 min, 10 min, and 30 min, respectively, before pelleting the exosomes at 100,000 g in a SW41Ti swinging bucket rotor for 3 hrs (Beckman). Supernatants (SNΔ) were collected, mixed with 5x Laemmli sample buffer, and incubated at 37°C for 10 min. For the exosomes (P100), samples were mixed with 1x Laemmli sample buffer and incubated at 37°C for 10 min. Then, the samples were analyzed by immunoblotting using anti-mouse Wnt3a, anti-GPR177 (17950-1-AP; Proteintech, IL, USA), or anti-TSG101 (ab125011, Abcam) antibodies.

### Dialysis

Purified Wnt3a (500 ng/mL) and 1% CHAPS were loaded into Slide-A-Lyzer Dialysis Device (10 kDa MWCO) with or without AFM or BSA. Dialysis against DMEM/F12 was performed for 48 hrs at 4 °C. During dialysis, the old dialysate (DMEM/F12) was discarded and replaced with fresh DMEM/F12 every 6 or 12 hrs. The samples were then transferred to new 1.5 mL LoBind tubes for further assays.

### Human organoids culture

Human normal stomach and colon tissue samples were obtained from patients who underwent gastrectomies, colectomies or colonoscopies. All samples were collected after obtaining informed consent from the patients before their participation in the study. The use of donor materials for research purposes was approved by the Institutional Review Board (IRB) of Yonsei University Health System (4-2012-0859 and 4-2017-0106). The tissue samples were minced, washed with ice-cold DPBS, and then incubated with a gentle cell dissociation reagent (Stemcell technologies) at room temperature for 30 min to release gastric glands or colonic crypts. Isolated gastric glands were washed with ice-cold DPBS, suspended in 25 μL of Matrigel, and seeded into 48-well plates. After 15 minutes for solidification of the Matrigel at 37°C, human stomach organoid culture medium was added to each well. Human stomach organoid culture medium contains advanced DMEM/F12 supplemented with 10 mM HEPES, 2 mM GlutaMax and 1x antibiotics-antimycotics (all from Gibco) with the following additional factors: 2% B-27 supplement (Gibco), 1 mM N-acetylcysteine (Sigma), 50 ng/ml EGF (Peprotech), 150 ng/ml noggin (Peprotech), 10% R-spondin1 CM (produced using HA-R-Spondin-1-Fc 293T cells, Trevigen), 200 ng/ml FGF10 (Peprotech), 10 nM gastrin-I (Sigma), 2 μM A-8301 (Tocris), and 50% Wnt3a CM for stomach organoids; 2% B-27 supplement, 1 mM N-acetylcysteine, 50 ng/ml EGF, 100 ng/ml noggin, 10% R-spondin1 CM, 10 nM gastrin-I, 500 nM A-8301, 10 μM SB202190 (Tocris), and 50% Wnt3a CM for colon organoids. The medium was replaced every 3 days, and the organoids were split every 7 days.

### Organoid growth assay

To assess the growth of organoids treated with each Wnt3a CM preparation, the organoids were digested with TrypLE (Gibco) at 37°C for 5–10 minutes until they were dissociated into single cells. Cells were washed with DMEM supplemented with 1% fetal bovine serum and 1% penicillin-streptomycin (Gibco) and plated on 24-well plates at a density of 5,000 cells/well in 40 μL of Matrigel. After 10 days of growth, the organoids were digested down to single cells using TrypLE and re-plated at the same density, while the cell numbers for each well were counted. The cumulative number of cells at each passage was calculated using the following equation: number of cells in the previous passage x the number of cells in the current passage / the number of cells plated in the current passage.

### Quantitative real-time RT-PCR

RNA extraction from organoids was conducted using the RNeasy Mini Kit (Qiagen) followed by cDNA synthesis with Superscript IV (Invitrogen) according to the manufacturer’s instructions. Then, cDNAs were amplified using SYBR Green Master mix (Applied Biosystems) on a StepOne Real-Time PCR system (Applied Biosystems). β-actin was used as an internal control and the expression level of each gene was normalized to the level of β-actin. The PCR primers used were as follows: 5’-AGGTCTGGTGTGTTGCTGAG-3’ and 5’-GTGAAGACGCTGA GGTTGGA-3’ for *LGR5* and 5’-GCTGCGCTTTGATAAGGTCC-3’ and 5’-GCTCATCTGAACCTCCTCT CTTT-3’ for *AXIN2*.

### Immunostaining

Organoids were fixed with 4% paraformaldehyde for 30 min at room temperature. Fixed organoids were embedded in Histogel (Thermo scientific) and further processed into paraffin blocks. Immunostaining was performed according to standard immunostaining protocols. Anti-Ki67 (1:200, Cell Signaling Technology #9449) was used, and fluorescent images were taken on an LSM 780 confocal microscope (Zeiss). Ki67-positive cells were counted and expressed as a percentage of DAPI-positive cells.

### Quantification and statistical analysis

The data were presented as means ± SEM. All statistical analyses were performed using GraphPad Prism 5 software. Data obtained from the luciferase reporter assays, MTT assays, immunoblots, quantitative real-time RT-PCR experiments, organoid growth assays, and immunostaining experiments were analyzed using the Student’s *t* test, ratio paired *t* test, one-way Analysis of Variance (ANOVA), two-way ANOVA or repeated measures two-way ANOVA techniques with post-hoc Bonferroni corrections for multiple comparisons. p-values < 0.05 were considered statistically significant. Additional details and the p-values for statistical significance are described in the figure legends.

## Acknowledgments

We would like to thank Dr. Hans Clevers for providing L-Wnt3a cells. This study was supported by grants from the National Research Foundation of Korea (NRF) funded by the Korean government (MSIT) (NRF-2017M3C7A1048090, NRF-2018R1A5A2025079 and NRF-2019R1A2C3002354 to C.H.K.; NRF-2018R1A6A1A03023718 to J.L.; 2021R1I1A1A01047269 to J.H.Y) and in part by a grant from Interpark Bio Convergence Corp., Seoul, Korea and a faculty research grant of Yonsei University College of Medicine (6-2017-0166).

## Author contributions

J.L. and C.H.K. conceived and designed experiments. J.H.Y., S.S.K., Y.O.J., J.C., J.H., and J.Y.L. performed the experiments. J.H.Y., S.S.K., J.C., I.K., J.L., Y.O.J., and C.H.K. analyzed the data. J.C. and T.K. contributed reagents or materials. J.H.Y., S.S.K., J.C., I.K., J.L., Y.O.J., and C.H.K. wrote the paper.

## Data availability

All data generated or analyzed for this study are included in the manuscript and supporting files; source data files have been provided for Figure 1-6.

---

**Figure 1 – Figure supplement 1.**
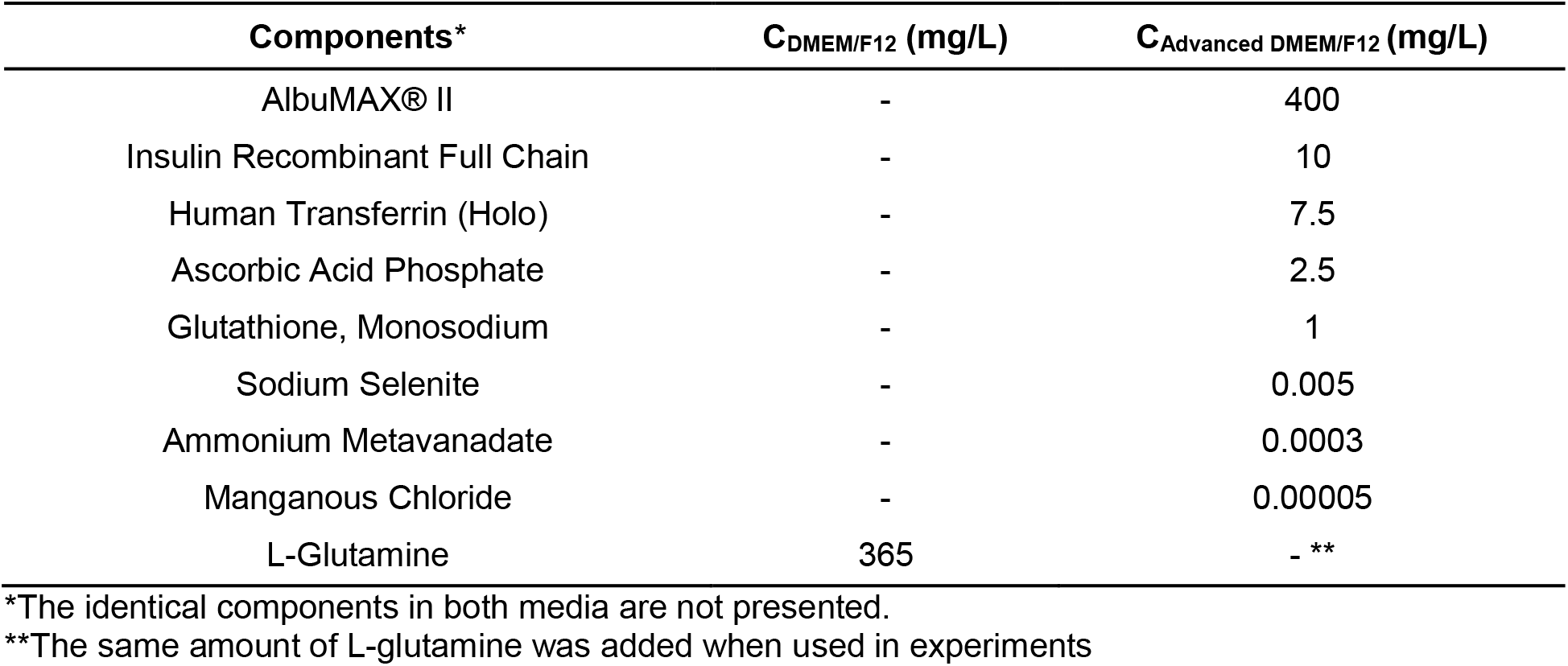
Formulation comparison between DMEM/F12 and AdDMEM/F12.

**Figure 1- Figure supplement 2.**
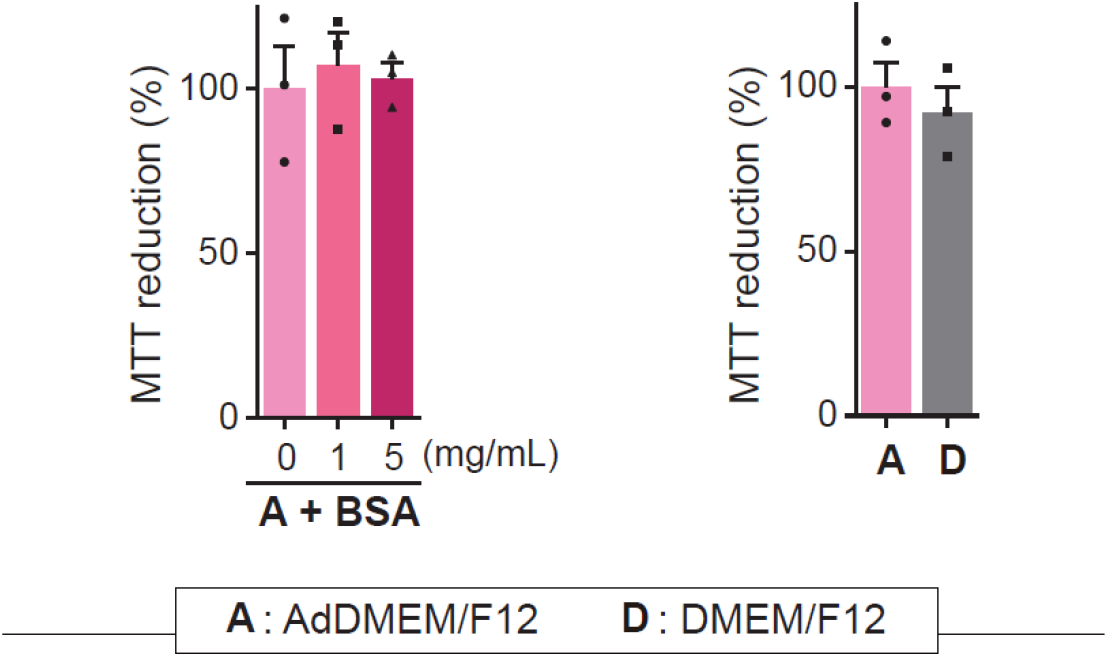
MTT assays in L-Wnt3a-AFM cells.

**Figure 3 – Figure supplement 1.**
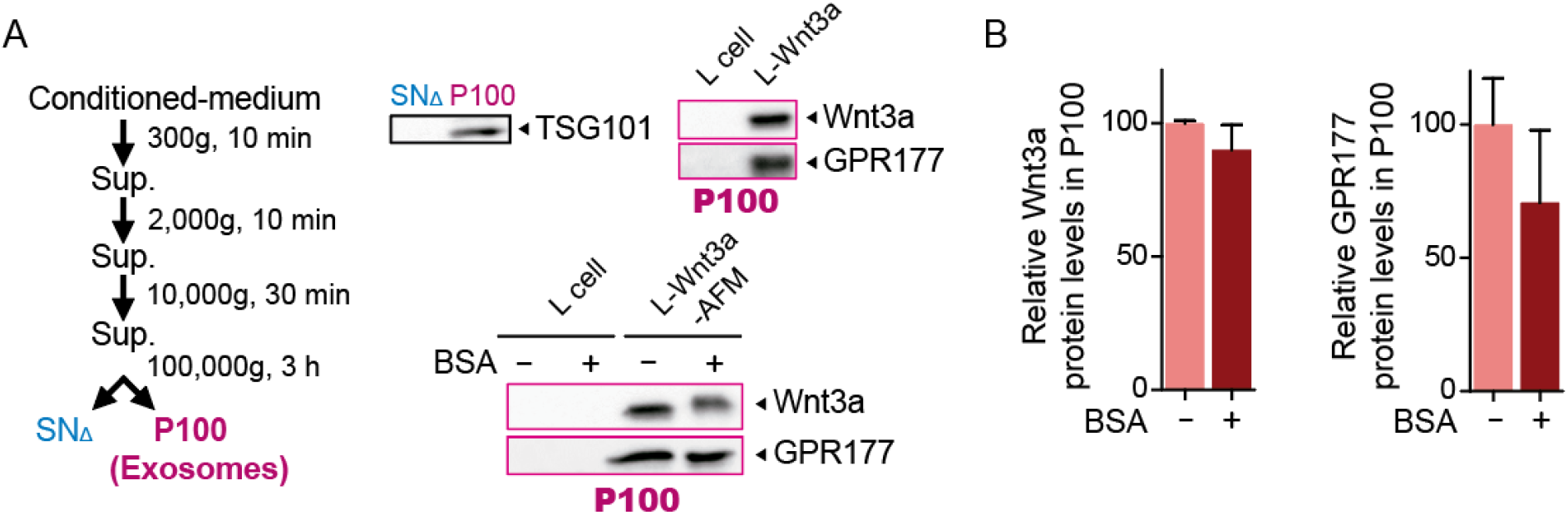
BSA does not affect Wnt3a secretion on exosomes.

**Figure 5 – Figure supplement 1.**
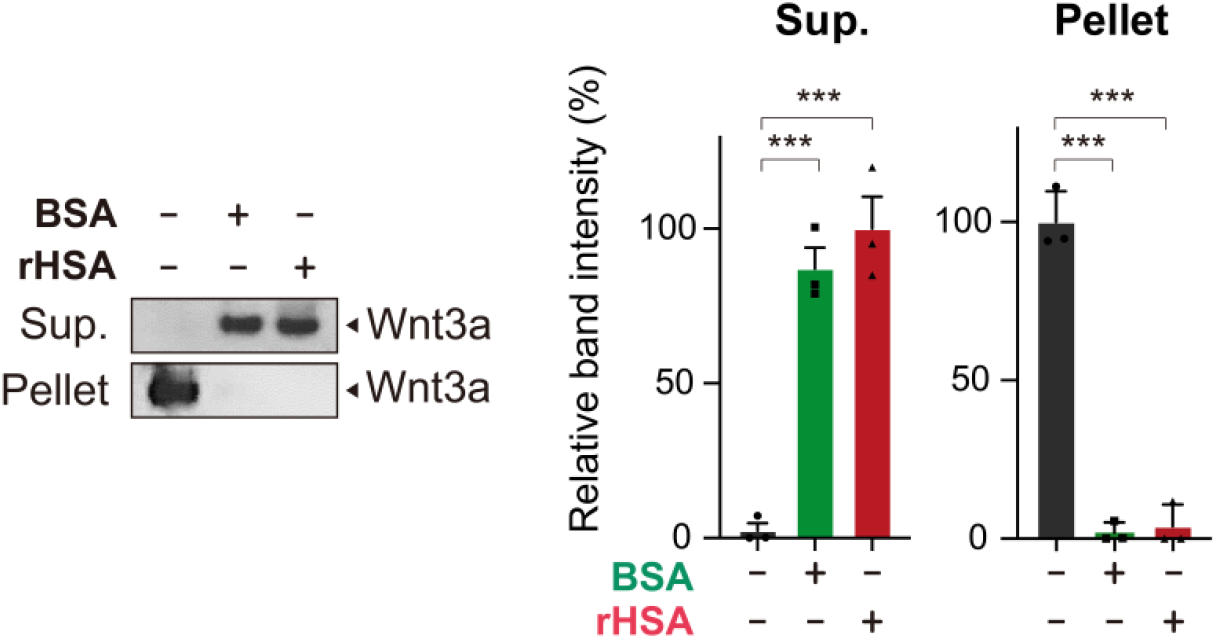
rHSA and BSA show similar effects on Wnt3a solubility.

**Figure 5 – Figure supplement 2.**
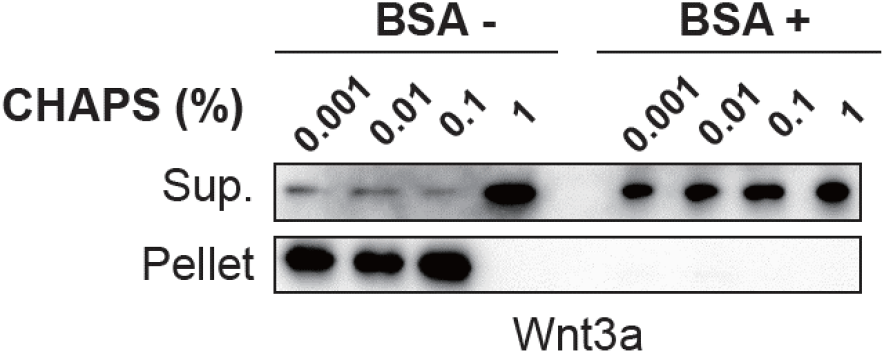
BSA efficiently solubilizes Wnt3a even in trace amounts of CHAPS.

**Figure 6 – Figure supplement 1.**
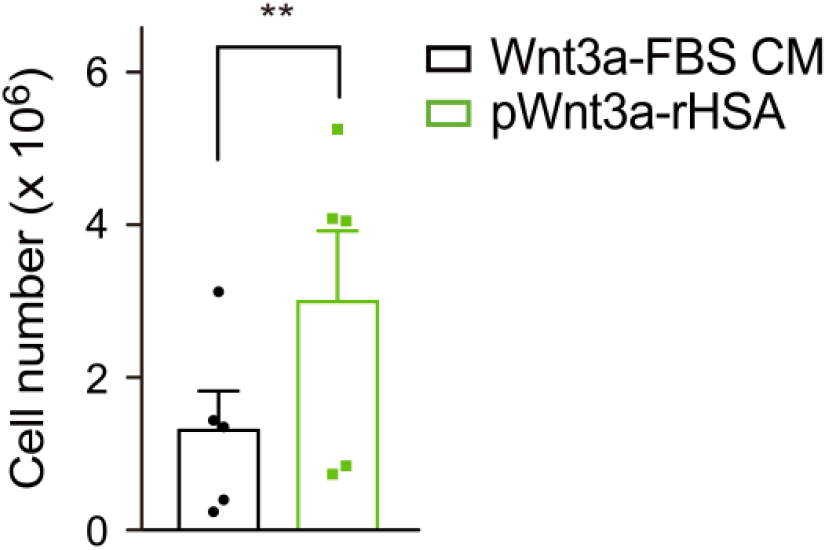
rHSA and BSA show similar effects on growth and expansion of human colon organoids.

**Source data – Figure 1-Source data 1. Raw data related to Figure 1B-1G; Original blots for Figure 1F**

**Source data – Figure 2-Source data 1. Raw data related to Figure 2A, B**

**Source data – Figure 3-Source data 1. Raw data related to Figure 3A-D; Original blots for Figure 3A-E**

**Source data – Figure 4-Souce data 1. Raw data related to Figure 4A-F**

**Source data – Figure 5-Source data 1. Raw data related to Figure 5B, 5C and 5E; Original blots for Figure 5B-F**

**Source data – Figure 6-Source data 1. Raw data related to Figure 6F-I and 6K; Original blots for Figure 6A, 6C-E**

